# Chromatin variation associated with liver metabolism is mediated by transposable elements

**DOI:** 10.1101/052571

**Authors:** Juan Du, Amy Leung, Candi Trac, Brian W. Parks, Aldons J. Lusis, Rama Natarajan, Dustin E. Schones

**Affiliations:** Department of Diabetes Complications and Metabolism, Beckman Research Institute, City of Hope, Duarte, California, United States of America; Irell & Manella Graduate School of Biological Sciences, City of Hope, Duarte, California, United States of America; Department of Nutritional Sciences, University of Wisconsin-Madison, Madison, Wisconsin, United States of America; Department of Medicine, University of California, Los Angeles, California, United States of America

**Keywords:** Chromatin accessibility, transposable element, transcription factor, DNA methylation, FAIRE-seq

## Abstract

**Background:** Functional regulatory regions in eukaryotic genomes are characterized by the disruption of nucleosomes leading to accessible chromatin. The modulation of chromatin accessibility is one of the key mediators of transcriptional regulation and variation in chromatin accessibility across individuals has been liked to complex traits and disease susceptibility. While mechanisms responsible for chromatin variation across individuals have been investigated, the overwhelming majority of chromatin variation remains unexplained. Furthermore, the processes through which the variation of chromatin accessibility contributes to phenotypic diversity remain poorly understood.

**Results:** We profiled chromatin accessibility in liver from seven strains of mice with phenotypic diversity in response to a high-fat/high-sucrose (HF/HS) diet and identified reproducible chromatin variation across the genome. We found that sites of variable chromatin accessibility were more likely to coincide with particular classes of transposable elements (TEs) than sites with common chromatin features. Evolutionarily younger long interspersed nuclear elements (LINEs) are particularly enriched for variable chromatin sites. These younger LINEs are enriched for binding sites of immune-associated transcription factors, whereas older LINEs are enriched for liver-specific transcription factors. Genomic region enrichment analysis indicates that variable chromatin sites at TEs contribute to liver metabolic pathways. Finally, we show that polymorphism of TEs and differential DNA methylation at TEs can both contribute to chromatin variation.

**Conclusions:** Our results demonstrate specific classes of TEs contribute to chromatin accessibility variation across strains of mice that display phenotypic diversity in response to a HF/HS diet. These results indicate that regulatory variation at TEs is an important contributor to phenotypic variation among populations.

## Background

Accessible (open) chromatin is a common feature of active regulatory regions in eukaryotic genomes [1, 2]. The cell type-specific accessibility of chromatin allows regulatory factors to bind the underlying DNA, leading to tightly regulated gene expression [1, 3, 4]. Accessible chromatin regions have been shown to be variable among different individuals [1, 5, 6], and these variable chromatin sites have been shown to be associated with complex traits and disease susceptibility [7]. However, the mechanisms underlying chromatin accessibility variation, and the processes through which this variation impacts phenotypic diversity, remain poorly understood.

Initial investigations into the relationship between variation of chromatin accessibility and genetic variation have begun to elucidate some principles. Examination of chromatin signatures in individuals with diverse ancestries revealed extensive variation in regulatory regions and evidence of heritability of these signatures [6]. Chromatin accessibility profiling in human lymphoblastoid cell lines revealed the association of chromatin accessibility signatures with genetic variants which are associated with the expression of nearby genes and potentially phenotypic diversity in humans [5, 8]. A study in erythroblasts from eight strains of inbred mice found that approximately 1/3 of variable open chromatin sites can be explained by single nucleotide variants and that these variants were associated with complex traits and disease [7]. While these pioneering studies have provided some insight into the drivers of chromatin variation, the majority of chromatin variation across the genome remains unexplained.

In addition to single nucleotide variants, transposable elements (TEs) constitute a major portion of genomic variation [9, 10]. Approximately 50% of the human genome and 40% of the mouse genome are derived from TEs [11, 12]. TEs can affect nearby gene activity, and have been linked to complex traits and diseases, including cancer and diabetes [13, 14]. Due to the deleterious nature of TE transposition, mammalian systems have a number of transcriptional and post-transcriptional mechanisms to silence TEs [15]. The major mechanisms responsible for the suppression of TE transposition are DNA methylation, histone methylation and RNA interference [15-17]. Most DNA methylation in mammals occurs within TE sequences in order to transcriptionally suppress TE activities [17, 18]. Indeed, in somatic cells, most TEs are epigenetically silenced by DNA methylation [19]. However, studies have shown that specific TEs can be derepressed in a tissue-specific manner [19-21]. For example, tissue-specific DNA hypomethylation within TEs has been shown to contribute to novel regulatory networks [19].

There is growing evidence that TEs have evolved for the benefit of the host, contributing to host genome expansion and genetic innovation [22]. TEs contribute to the regulation of gene expression, by functioning as distal enhancers, alternative promoters or alternative splicing signals [19, 20, 23, 24]. Although TE-associated accessible chromatin sequences are less conserved among different species, the lack of repression of TEs have been associated with the transcription of nearby genes in a tissue-specific manner, indicating their functional relevance [25, 26]. Previous studies have identified binding sites of many transcription factors (TFs) within specific TE sequences [26, 27]. Analysis of TE-associated TF binding sites in different species has further suggested that the expansion of the mammalian TF binding repertoire has been mediated by TE transposition [24, 27]. Given the prevalence of TE sequences and their potential regulatory functions, we hypothesized that variation of TE regulation in strains of mice is a major cause of chromatin accessibility variation, and can further contribute to regulatory and phenotypic diversity.

To study the roles of TEs in chromatin accessibility variation, we chose seven strains of inbred mice that have differential response to a “western” high-fat / high-sucrose (HF/HS) diet [31] and performed genome-wide chromatin accessibility profiling in liver tissue using FAIRE-seq [32]. Given that TEs are typically repressed in somatic cells [15, 17], we expected that TEs would be less enriched at accessible chromatin. Interestingly, 37% of the variable chromatin regions are associated with TEs. Furthermore, the TE-associated regions of chromatin variation among different strains contribute to liver transcription regulation and metabolic pathways. Taken together, our study indicates that TEs are a major contributor to chromatin accessibility variation among different inbred strains and further contribute to phenotypic diversity.

## Results

### Chromatin accessibility variation observed in livers of mice with differential phenotypes

Previous studies have reported strain-specific heterogeneity in physiological response to HF/HS diet feeding [31, 33]. In this study, we chose seven commonly used inbred strains of mice: A/J, AKR/J, BALB/cJ, C57BL/6J, C3H/HeJ, CBA/J and DBA/2J. These strains display diverse body fat percentage change after 8 weeks of HF/HS feeding, ranging from an average increase of 70% (BALB/cJ) to over 200% (C57BL/6J) (Additional file 1: Figure S1a) [31]. We also observed significant variation of liver phenotypic markers, including liver triglyceride content (Fig. 1a; Additional file 1: Figure S1b) [34], as expected given the important metabolic processes that occur in the liver [35].

**Fig. 1.**
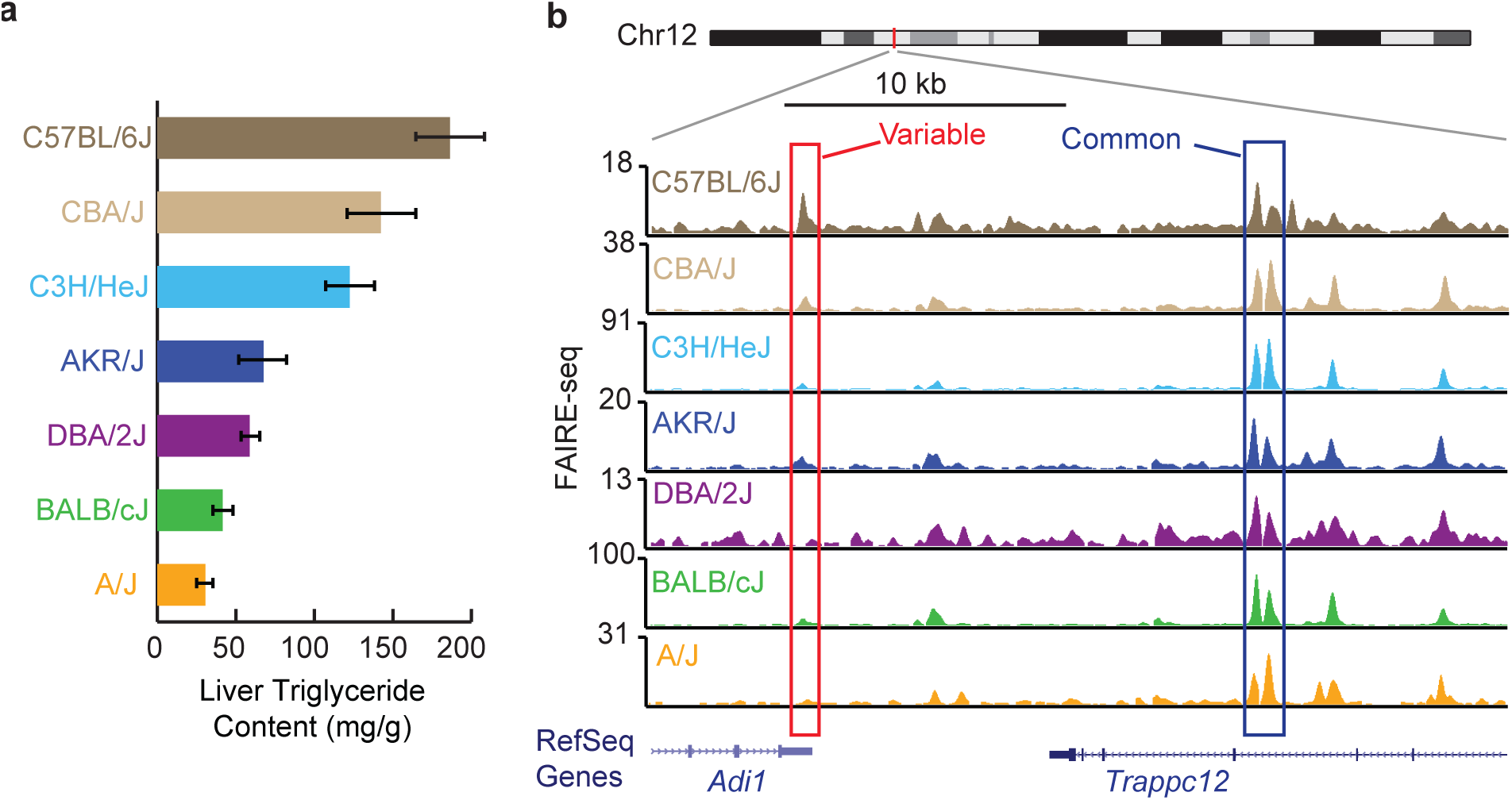
Chromatin variability across inbred strains of mice. (**a**) Liver triglyceride content of mice after 8 weeks of HF/HS feeding. The error bars represent the standard deviation of the measurements on 4 mice for each strain (adapted from Hui *et al.*, Elife, 2015. 4). (**b**) Genome browser view of variable and common chromatin sites.

To profile chromatin accessibility at a genome-wide level, we performed FAIRE-seq [32] in livers from these strains after 8 weeks of HF/HS feeding (two biological replicates for each strain). In order to mitigate alignment biases, we created strain-specific pseudo-genomes using known single nucleotide polymorphisms (SNPs) [36], as described previously [7]. We identified an average of 29,752 reproducible accessible chromatin sites in each individual strain (Additional file 2: Table S1). All together, we identified a union set of 50,775 open chromatin sites across the seven strains. To characterize variation of chromatin accessibility among different strains, we compared quantile-normalized read counts at the union set of sites using the DESeq package [37], as described previously [7]. We ranked sites by their adjusted *p*-value of variation (Additional file 1: Figure S2a) and selected the top 5% as the most variable set (2,539 sites; adjusted *p* < 1.21e-9). Similarly, we classified the bottom 5% of sites as common sites. Variable chromatin sites display dynamic patterns of chromatin accessibility across different strains (Additional file 1: Figure S2b). Examples of variable and common chromatin sites are shown in Fig. 1b.

Given that genetic variation in the form of SNPs has been shown to contribute to chromatin variation in mouse erythroblasts [7], we first tested if SNPs are associated with chromatin variation in the liver. We found that 30% (764/2539) of the most variable chromatin sites have underlying SNPs that are associated with chromatin variation among the seven strains (Additional file 1: Figure S3). This result is consistent with a previous study using erythroblasts from eight strains of inbred mice [7]. While providing a genetic explanation for ~1/3 of chromatin variation, this leaves the majority of chromatin variation among different inbred strains unexplained. We next turned our focus to identifying other potential genetics mechanisms contributing to chromatin accessibility variation.

### Chromatin variability at TEs across inbred strains

Previous studies have shown widespread contribution of TEs to regulatory networks in mammalian genomes [26]. We therefore reasoned that TEs could influence chromatin accessibility variation among inbred strains. Given that TEs are typically repressed/silenced in somatic cells [15, 17], we expected that TEs would be less enriched at sites of chromatin accessibility compared to random sites. To test this, we examined the prevalence of TEs in all accessible chromatin sites utilizing the RepeatMasker [38] annotation of TEs. As expected, sites of accessible chromatin are less likely to overlap instances of four classes of TEs (LINEs (long interspersed nuclear elements), SINEs (short interspersed nuclear elements), LTRs (long terminal repeats) and DNA transposons) compared to random sites in the genome (34% vs. 54%, *p* < 2.2 × e^−16^, Fisher’s exact test; Additional file 1: Figure S4a, b). These percentages are comparable to a previous study using DNase I hypersensitivity data sets from human tissues [26].

Interestingly, although TE sequences generally display less accessible chromatin, there are more TEs observed at variable chromatin sites than compared to common chromatin sites (37% vs 32%, *p* = 0.00105, Fisher’s exact test; Additional file 1: Figure S4c, d). Furthermore, two specific classes of TEs, LINEs and LTRs, are significantly more enriched at variable chromatin sites compared to common chromatin sites (Fig. 2a, b; LINE: *p* = 3.729 × e^−13^; LTR: *p* = 4.536 × e^−13^, Fisher’s exact test). As an example, the variable chromatin site at the *Adi1* locus in Fig. 1b coincides with a LTR (Additional file 1: Figure S4e). In contrast, SINEs are more enriched at common chromatin sites compared to variable sites (Fig. 2c, *p* = 0.00017, Fisher’s exact test). DNA transposons are not enriched at either variable or common sites (Additional file 1: Figure S5a, *p* = 0.37, Fisher’s exact test). Similar analysis of accessible chromatin profiles [30, 39] from three strains of mice (A/J, C57BL/6J and DBA/2J) fed a standard chow diet revealed a similar percentage of variable chromatin sites overlapping with TEs (42%, 876/2095), indicating that this is a general phenomenon across different strains independent of diet.

**Fig. 2.**
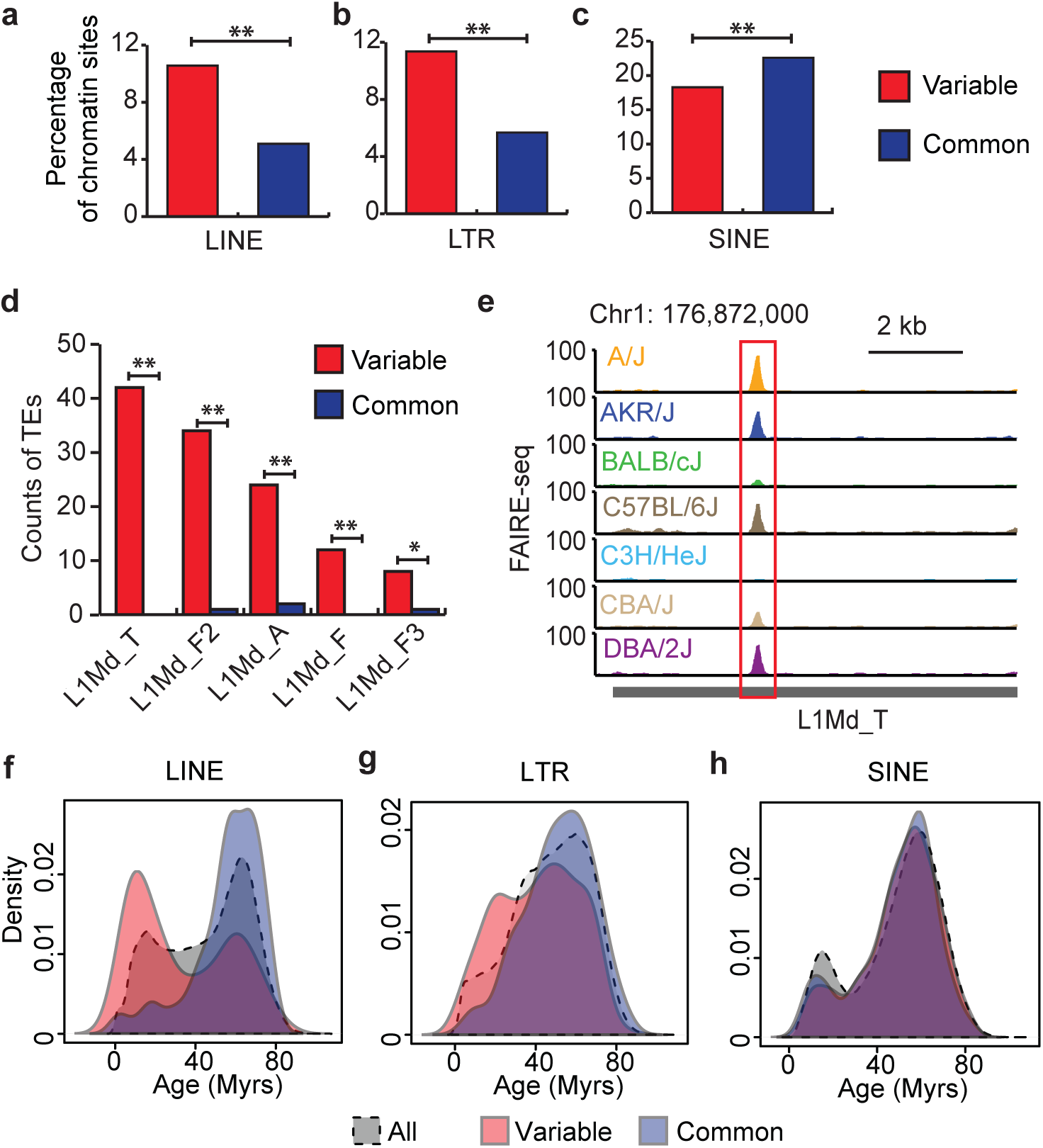
Specific subfamilies of TEs are enriched for variable chromatin sites. (**a** – **c**) Percentage of variable and common chromatin sites overlapping (**a**) LINEs, (**b**) LTRs and (**c**) or SINEs (** *p* < 0.001, Fisher’s exact test). (**d**) Numbers of variable (red) or common (blue) chromatin sites overlapping certain families of TEs (\**p* < 0.05, ** *p* < 0.001, Fisher’s exact test). (**e**) Genome browser view of a variable chromatin site in an L1Md_T element. (**f** – **h**) Age distribution of all TEs, TEs at variable or common chromatin sites for different classes of TEs, including (**f**) LINEs, (**g**) LTRs and (**h**) SINEs. Myrs: million years.

### Variable chromatin sites are enriched at evolutionarily younger LINEs

Given the evidence that specific subfamilies of TEs can play specific regulatory roles [27, 40], we next investigated if variable chromatin sites are enriched for specific families of TEs. Similar to previous analysis [24], we used the RepeatMasker [38] annotation of TE families and subfamilies and tabulated the occurrences of TEs from each subfamily at variable or common chromatin sites (Additional file 3: Table S2). Intriguingly, we found that several L1Md subfamilies are significantly enriched at the variable sites compared to common sites (Fig. 2d, e). These L1Md subfamilies of TEs are evolutionarily younger compared to other TEs. The average age of L1Md_T, L1Md_F2, L1Md_A, L1Md_F and L1Md_F3 are 8.27, 15.06, 8.05, 30.29 and 12.05 million years, respectively (see Methods) [27, 38]. RNA transcripts of L1Md elements have been detected in mouse liver, indicating these TEs can be somatically derepressed [41, 42]. While the function of these L1Md elements is largely unknown, the accessible chromatin signature we observed at these elements indicates that they may play a regulatory role in mouse liver.

Given that sites of variable chromatin are enriched for evolutionarily younger families of LINE elements compared to common sites, we next asked whether TEs at variable sites are in general evolutionarily younger than those at common sites. We again separated TEs into four classes (SINEs, LINEs, LTRs and DNA transposons) and plotted the distribution of the evolutionary age of all TEs as well as those at variable and common sites separately in each of the classes (Fig. 2f-h; Additional file 1: Figure S5b). Strikingly, we found that LINEs at variable chromatin sites display a bimodal distribution for age, with one subgroup evolutionarily younger compared to all LINEs (Fig. 2f). In contrast, LINEs that overlap common chromatin sites are in general evolutionarily older (Fig. 2f, *p* < 2.2 × e^−16^, Wilcoxon rank sum test). We observed a similar, albeit less dramatic, trend for LTR elements (Fig. 2g, *p* = 2.659 × e^−7^, Wilcoxon rank sum test). However, for SINE elements, there was no significant age difference observed between variable and common chromatin sites (Fig. 2h, p = 0.22, Wilcoxon rank sum test). DNA transposons that overlap variable chromatin show slight enrichment at older elements (Additional file 1: Figure S5b, *p* = 0.049, Wilcoxon rank sum test). However, DNA transposons contribute to a much smaller population of variable chromatin sites as compared to other classes of TEs (Fig. 2a-c, Additional file 1: Figure S5a). These results indicate that younger TEs, especially LINEs, are regions of increased regulatory differences across strains of mice and therefore may be involved in more recent adaptations of regulatory networks.

### Increased chromatin accessibility at younger LINEs

In order to understand the regulatory roles of younger LINEs, we examined the chromatin accessibility differences at all LINEs ranked by their evolutionary age (Fig. 3a). To better examine the coverage of all mappable reads from FAIRE-seq data at repetitive elements, we mapped FAIRE-seq reads to the mouse genome using bowtie2 [43], as described previously [44] (see Methods). To examine chromatin accessibility differences among younger and older LINEs, we ranked all RepeatMasker annotated LINEs by their evolutionary age, and then plotted FAIRE-seq read counts upstream and downstream of the annotated 5’ start and 3’ end of all LINEs (Fig. 3a). Interestingly, we found enriched chromatin accessibility at younger LINEs compared to older LINEs (Fig. 3a). The length of LINE elements in the genome is variable, with vast majority of LINEs being truncated [45]. To further examine the location of enriched chromatin accessibility within younger LINEs, we separated the LINEs into different groups based on their size and surveyed the FAIRE-seq at the LINEs and flanking regions (Fig. 3b). Consistent with the heatmap (Fig. 3a), younger LINEs have higher chromatin accessibility compared to older LINEs, regardless of size groups (Fig. 3b). We also found that longer LINEs have more enriched chromatin accessibility compared to shorter ones (Fig. 3b). Chromatin accessibly in another strain of mice, A/J, reveals a similar trend (Additional file 1: Figure S6). It has previously been shown that intact, longer, LINEs can be transcribed [20]. However, we did not detect RNA transcripts at LINEs with RNA-seq (Fig. 3c). One possibility for this is that TE transcripts are subjected to post-transcriptional suppression that affects RNA stability [15]. All together, these results indicate that a group of evolutionarily younger LINEs have regulatory functions in mouse liver, while not producing stable transcripts.

**Fig. 3.**
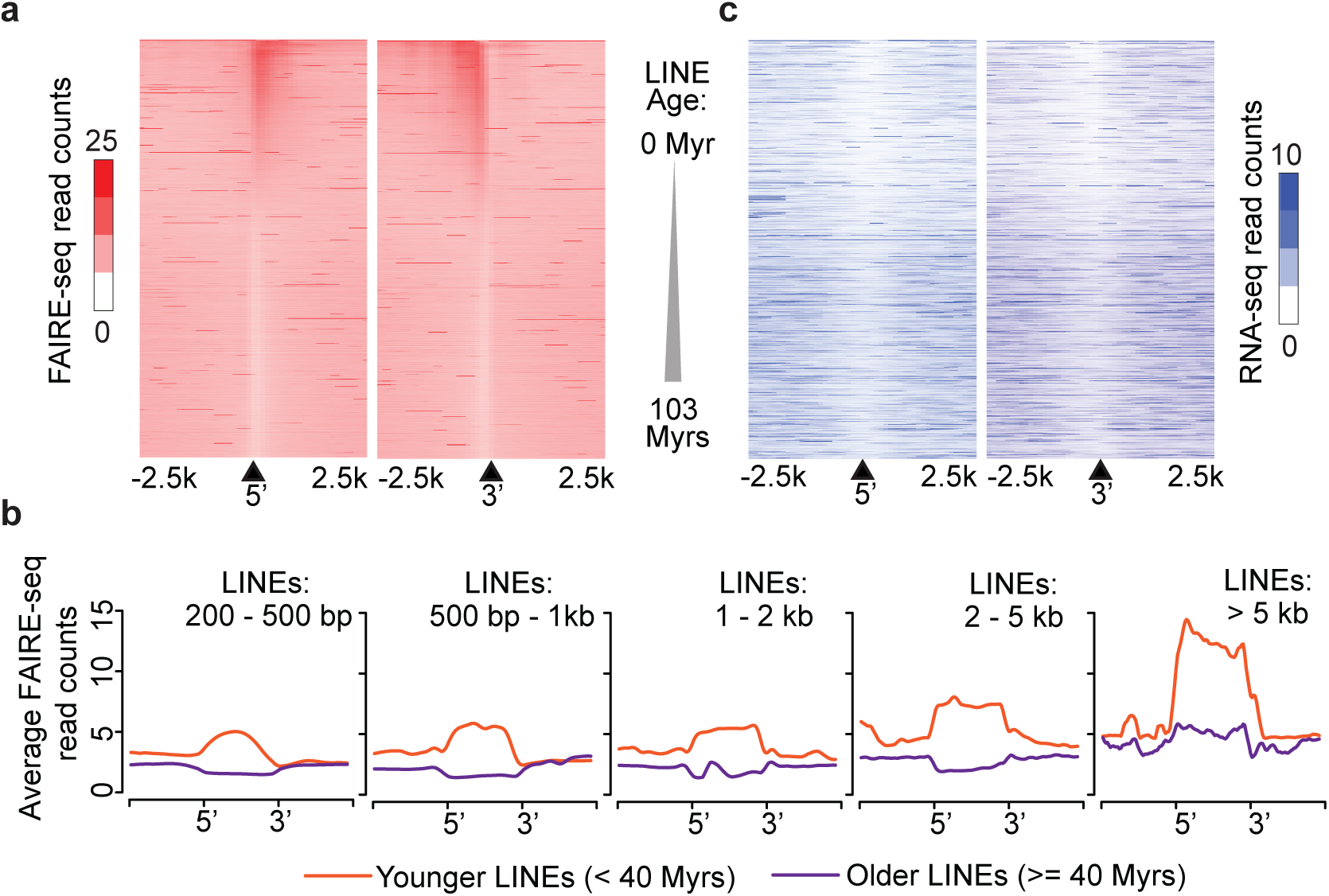
Differential chromatin accessibility profiles at younger and older LINEs. Heatmap showing FAIRE-seq read counts from C57BL/6J mice liver surrounding the 5’ and 3’ regions of LINEs sorted by their evolutionary age. Black triangles denote the 5’ (left) or the 3’ (right) end of LINEs, with counts extending +/− 2,500 bp upstream and downstream. (**b**) Aggregate plots of average FAIRE-seq read counts upstream, downstream and within LINEs, stratified by size of LINEs. (**c**) Heatmap showing RNA-seq read counts from C57BL/6J mice liver surrounding the 5’ and 3’ regions of LINEs sorted by their evolutionary age. Myrs: million years.

### Differential transcription factor binding repertoire contributed by younger and older LINEs

TEs have been shown to contain transcription factor (TF) binding sites, and contribute to the evolution of the mammalian TF binding repertoire [24, 27, 46]. We examined the potential regulatory roles of LINEs by scanning for binding sites of known TFs in LINE-associated variable chromatin sites (see Methods). We scanned for enriched known motifs in variable chromatin sites containing younger or older LINEs. We found that different TF binding motifs are enriched at sites overlapping older LINEs compared to those overlapping younger LINEs (Fig. 4a). The motif for HNF4α, a liver TF, is the top enriched motif in variable chromatin sites containing older LINEs (Fig. 4a). HNF4α ChIP-seq data from C57BL/6J liver [47] also validated the enrichment of HNF4α binding at older LINEs compared to younger ones (Fig. 4b; *p* < 2.2 × e^−16^, Wilcoxon rank sum test). In addition, the binding motif for another liver TF, C/EBPα, is also enriched at variable sites containing older LINEs (Fig. 4a). Notably, 59% (67/114) of variable chromatin sites overlapping older LINEs are bound by the two liver TFs, HNF4α and/or C/EBPα. To serve as a control, we searched for the sites that are bound by CTCF [4], a non-liver-specific TFs. Compared to HNF4α and/or C/EBPα, we found only 11% (12/114) of variable chromatin sites overlapping older LINEs to be bound by CTCF, indicating the important role of older LINEs in liver-specific transcription regulation (Fig. 4c, *p* = 7.487 × e^−15^, Fisher’s exact test).

**Fig. 4.**
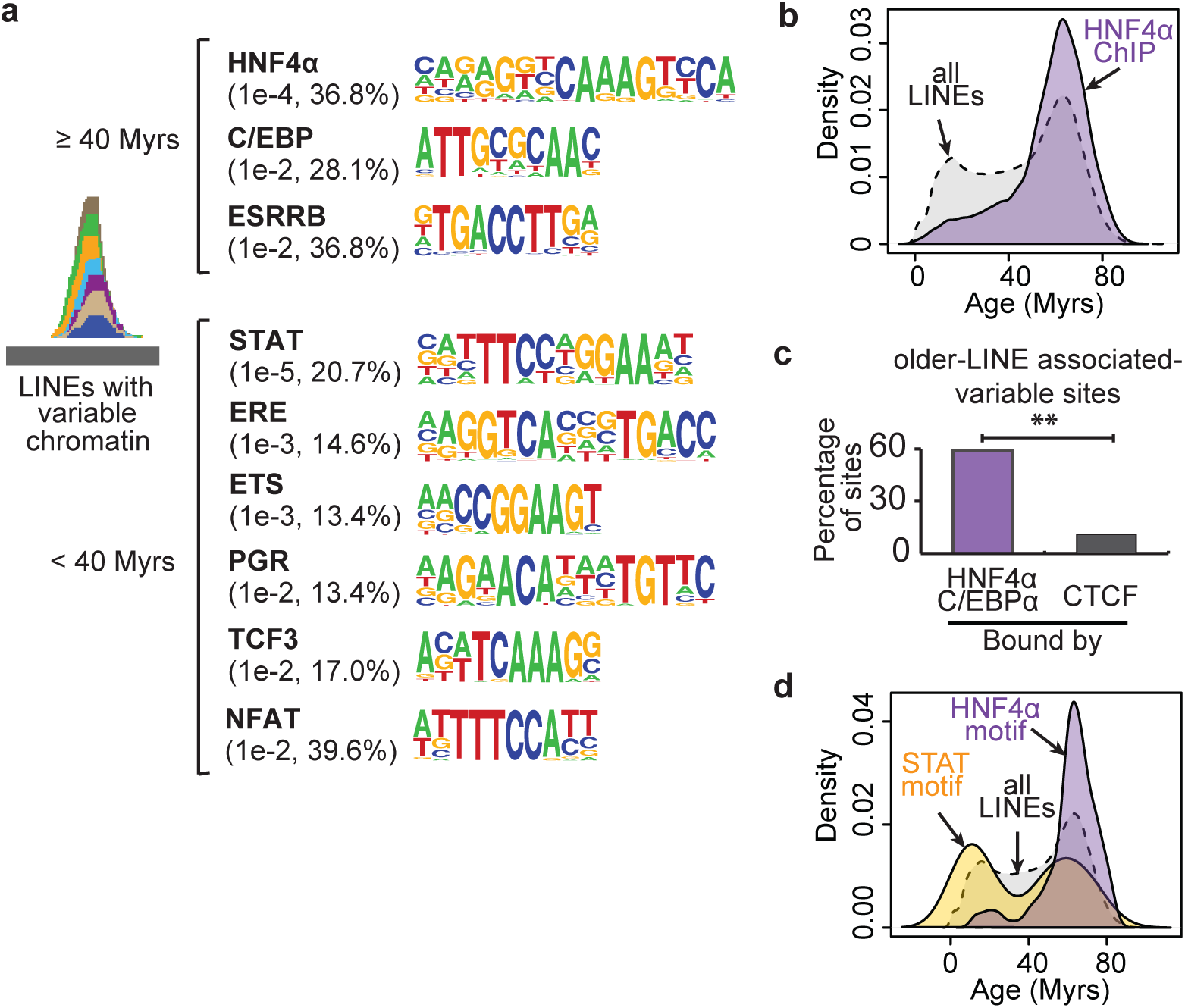
Specific TFs bind to younger and older LINEs. Top known motifs found in variable chromatin sites overlapping older (top) or younger (bottom) LINEs. Numbers in brackets represent p-values of enrichment of motif occurrence in given sequences compared to background, and the percentage of variable chromatin sites harboring the motif. (**b**) Age distribution of all LINEs and LINEs containing HNF4~ ChIP-seq peaks. (**c**) Percentage of older-LINE associated-variable sites bound by liver TFs and CTCF (** *p* < 0.001, Fisher’s exact test). (**d**) Age distribution of all LINEs, LINEs containing STAT or HNF4~ motif within accessible chromatin regions. Myrs: million years.

Intriguingly, variable chromatin sites containing younger LINEs are most enriched for the binding motif of STAT proteins (Fig. 4a), which have been shown to play an important role in response to inflammation in liver [48]. In addition, we noticed that several other enriched motifs contain a half GAS motif (TTC or GAA), which STATs can also bind to [49, 50]. To further investigate the presence of specific TF binding at specific LINEs, we used the occurrence of motifs at accessible chromatin sites in C57BL/6J mice liver to predict binding [51]. Of the predicted HNF4α binding sites, 87% (7209/8296) have HNF4α ChIP-seq peaks in mouse liver [47]. Using these predicted binding sites, we found that compared to older LINEs that are enriched for HNF4α binding sites, younger LINEs at accessible chromatin regions are enriched for STAT binding sites (Fig. 4d; *p* = 2.318 × e^−9^, Wilcoxon rank sum test). These results indicate that younger LINEs may play a role in the STAT-mediated immune response in the liver. Given that these younger LINEs are enriched at variable chromatin sites, differential STAT binding may also contribute to the phenotypic variation among different strains. Importantly, these results indicate that specific LINE elements of different evolutionary age have contributed unique elements to regulatory networks.

### TE-associated variable chromatin sites contribute to liver metabolic pathways

To further investigate the contribution of variable chromatin at TEs to phenotypic diversity among strains (Fig. 1a; Additional file 1: Figure S1), we used the Genomic Regions Enrichment of Annotations Tool (GREAT) [52] to identify enriched biological functions of accessible chromatin sites overlapping TEs. We found that variable chromatin sites with TEs are enriched in liver metabolic pathways, including gluconeogenesis, insulin secretion and lipid storage (Fig. 5a). More specifically, the variable chromatin sites containing younger LINEs are enriched in the negative regulation of gluconeogenesis (Additional file 4: Table S3). Interestingly, a previous study has shown that STAT3 regulates expression of gluconeogenic genes in liver [53], indicating that younger LINEs harboring STAT binding motifs are playing a role in regulating gluconeogenesis in liver. In contrast, variable chromatin sites at unique sequences (without TE or other repeats) of the genome are only enriched for filopodium assembly and antigen processing pathways (Fig. 5a). In addition, variable chromatin sites overlapping other types of repeats show no enrichment for biological functions, and comprise only a small percentage of variable chromatin sites. To serve as a control, we also searched for enriched biological processes in common chromatin sites with or without TEs. Not surprisingly, both groups of common chromatin sites are enriched for liver metabolic processes, including triglyceride metabolic process and cellular response to oxidative stress (Additional file 4: Table S3), indicating that these liver metabolic pathways are conserved and tightly regulated in all the strains. Taken together, these results suggest that the variation of chromatin accessibility among different strains is associated with liver metabolic pathways through specific TE sequences.

**Fig. 5.**
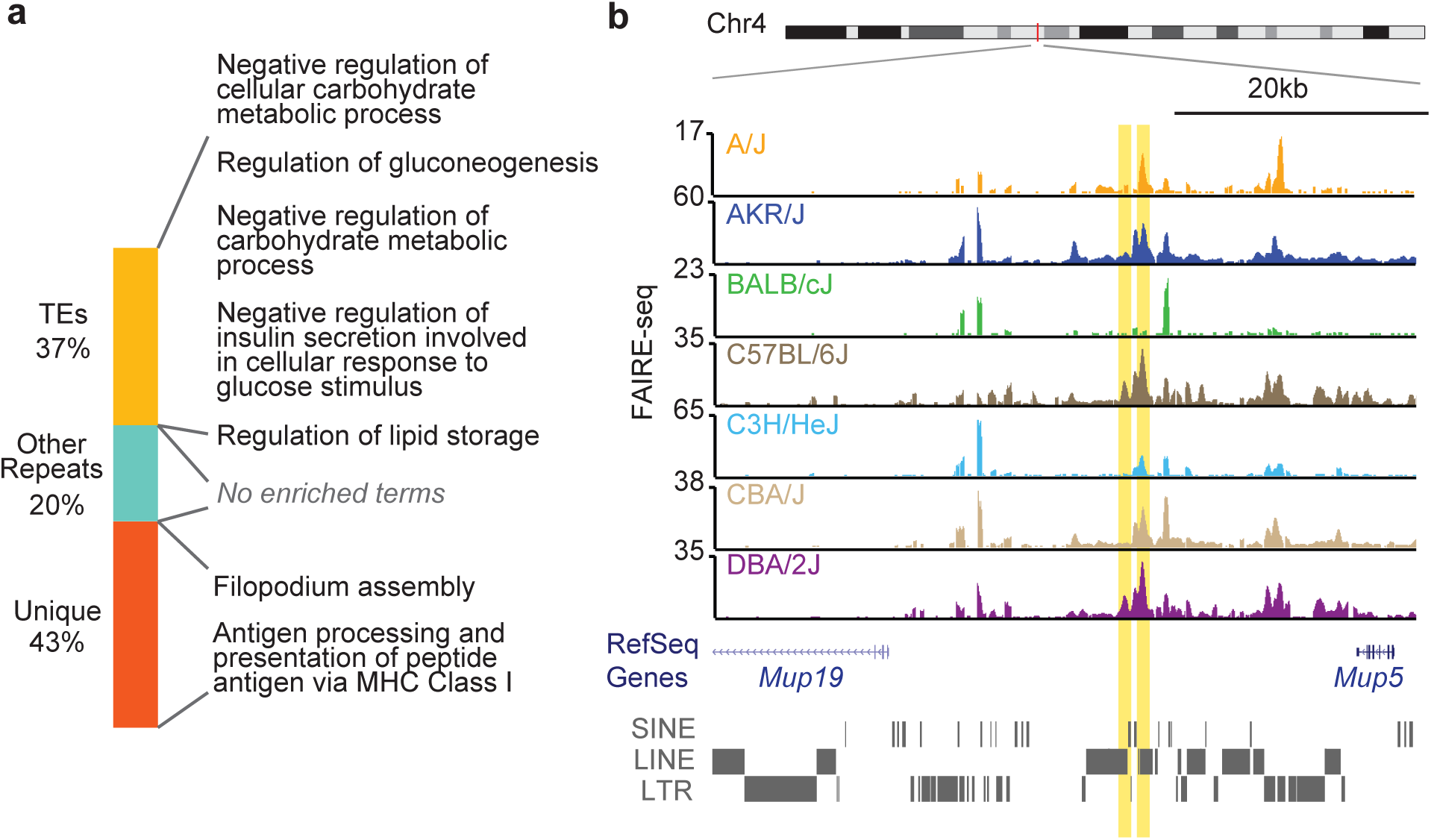
TE-associated chromatin variation contributes to liver phenotype. Enriched biological process of variable chromatin sites. Genomic coordinates of accessible chromatin sites were used as input for Genomic Regions Enrichment of Annotations Tool (GREAT) analysis (see Methods). (**b**) Genome browser view of TE-associated variable chromatin sites within *Mup* locus.

As an example, several TE-associated variable chromatin sites are found in the major urinary protein (MUP) gene locus. Two LINE-associated variable sites proximal to *Mup19* and *Mup5* are shown in Fig. 5b. MUP family proteins are expressed mainly in the liver and bind to small lipophilic molecules, including fatty acids [54]. MUPs have been shown to play important roles in glucose and lipid metabolism, and are highly polymorphic in mice [54-56]. Our results here suggest that TEs are involved in this polymorphic feature of MUPs in the mouse genome.

### TE polymorphic variants contribute to regulatory variation across inbred strains

Given the widespread contribution of TEs to regulatory networks, we were further interested in characterizing the potential mechanisms responsible for TE-driven regulatory variation among different strains. One possible mechanism whereby TEs could contribute to chromatin accessibility variation is TE polymorphism – where a TE is present in one genome and not in another (Fig. 6a). A previous study has characterized TE polymorphism across 18 strains of mice, including the seven strains in our study [10].

**Fig. 6.**
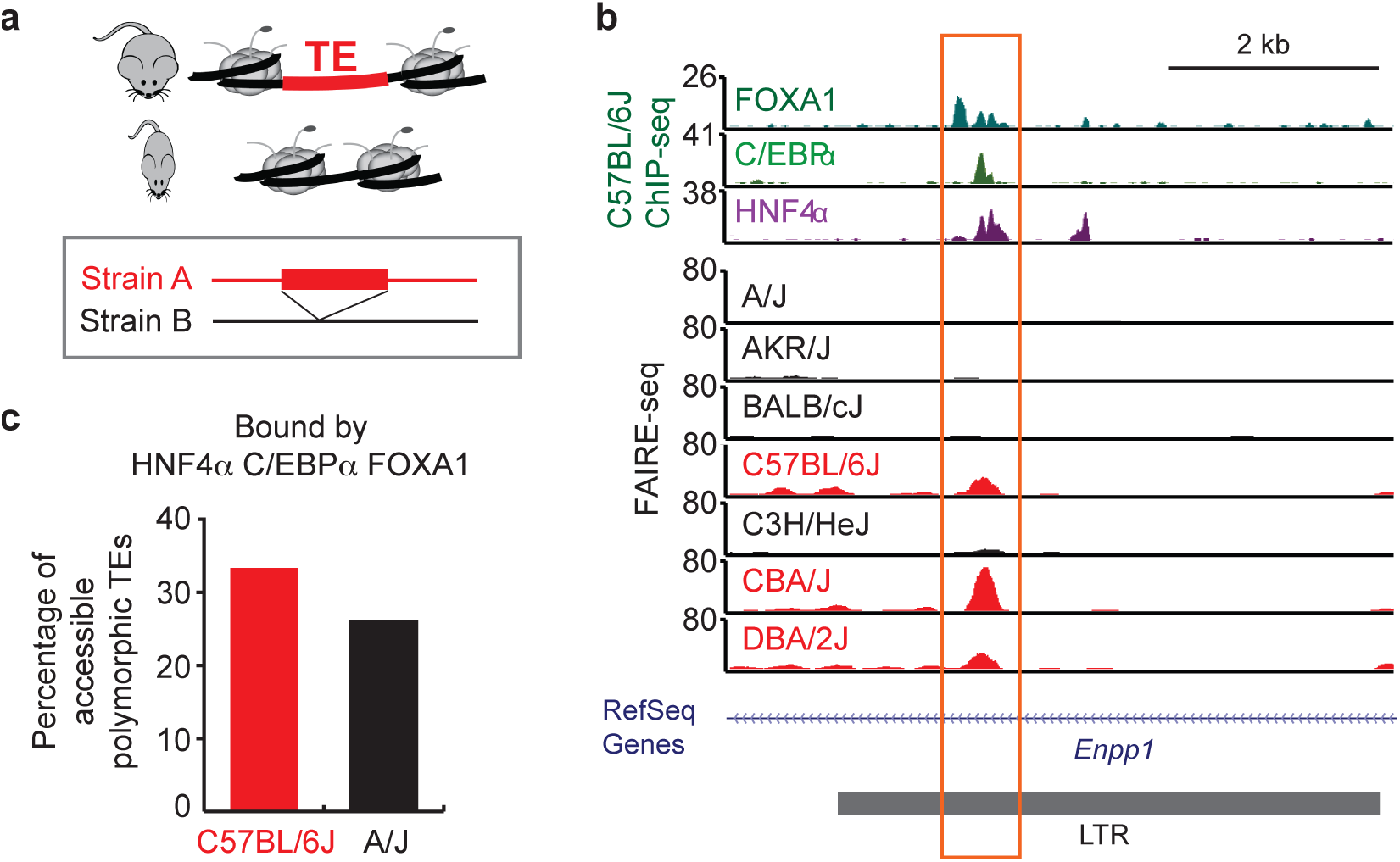
Contribution of TE polymorphism to chromatin variation. Model of TEs contributing to chromatin variation through TE polymorphism. (**b**) Genome browser view of a polymorphic TE variant with variable chromatin accessibility. Strains with FAIRE-seq tracks colored in red contain the LTR element, while the strains in black do not have the LTR element. (**c**) Percentage of polymorphic TE-associated variable chromatin sites shown to be bound by liver TFs in C57BL/6J or A/J mice livers.

Fig. 6b shows an example of a polymorphic LTR variant associated with chromatin variation. The LTR element present in C57BL/6J, CBA/J and DBA/2J genomes [10] contains a strain-specific accessible chromatin region. Interestingly, the accessible chromatin site within the LTR also shows evidence of binding by liver TFs, including HNF4α, C/EBPα, and FOXA1 (Fig. 6b). This region is within the intron of the *Enpp1* gene, which encodes a pyrophosphatase, and has been shown to be related to type 2 diabetes [57]. These results indicate that polymorphic TE-associated chromatin sites may play a strain-specific regulatory role for *Enpp1*. While we found that only 6% (59/934) of the TEs that overlap with variable chromatin sites are polymorphic amongst these strains, this is likely an underestimate given the difficulty in genome assembly at repetitive regions of the genome [10]. Previous work has shown that polymorphic TE variants that survived purifying selection rarely result in gene expression changes [10]. However, we found approximately 30% of polymorphic TE sites are bound by liver TFs (Fig. 6c), suggesting that these polymorphic TE-associated variable chromatin sites are playing regulatory roles.

### Differential DNA methylation at TEs contributes to regulatory variation across inbred strains

It has been previously demonstrated that TEs are subject to regulation through epigenetic mechanisms, including DNA methylation and histone modifications [17]. In human somatic cells, DNA hypomethylation has been found within specific TE subfamilies that are associated with enhancer marks [19]. We therefore reasoned that TEs with differential chromatin accessibility not classified as polymorphic could be differentially regulated through epigenetic mechanisms, such as DNA methylation (Fig. 7a). An example of a TE-containing variable chromatin locus with negatively associated CpG methylation levels is shown in Fig. 7b,c. Interestingly, strain-specific (A/J vs C57BL/6J) binding of liver TFs [47] indicates that this region is differentially bound by liver TFs as well (Fig. 7b). To examine the contribution of differential methylation at TEs to chromatin variation across the genome, we utilized reduced representation bisulfite sequencing (RRBS) data from liver tissue of the same strains of mice [58]. Interestingly, variable chromatin sites at TEs have a greater degree of DNA methylation variation across strains as compared to variable chromatin sites at other regions (Fig. 7d), emphasizing that differential DNA methylation at TEs is a major contributor to chromatin variation. These results indicate that differential epigenetic suppression of TEs contributes to regulatory variation in livers of different inbred mouse strains.

**Fig. 7.**
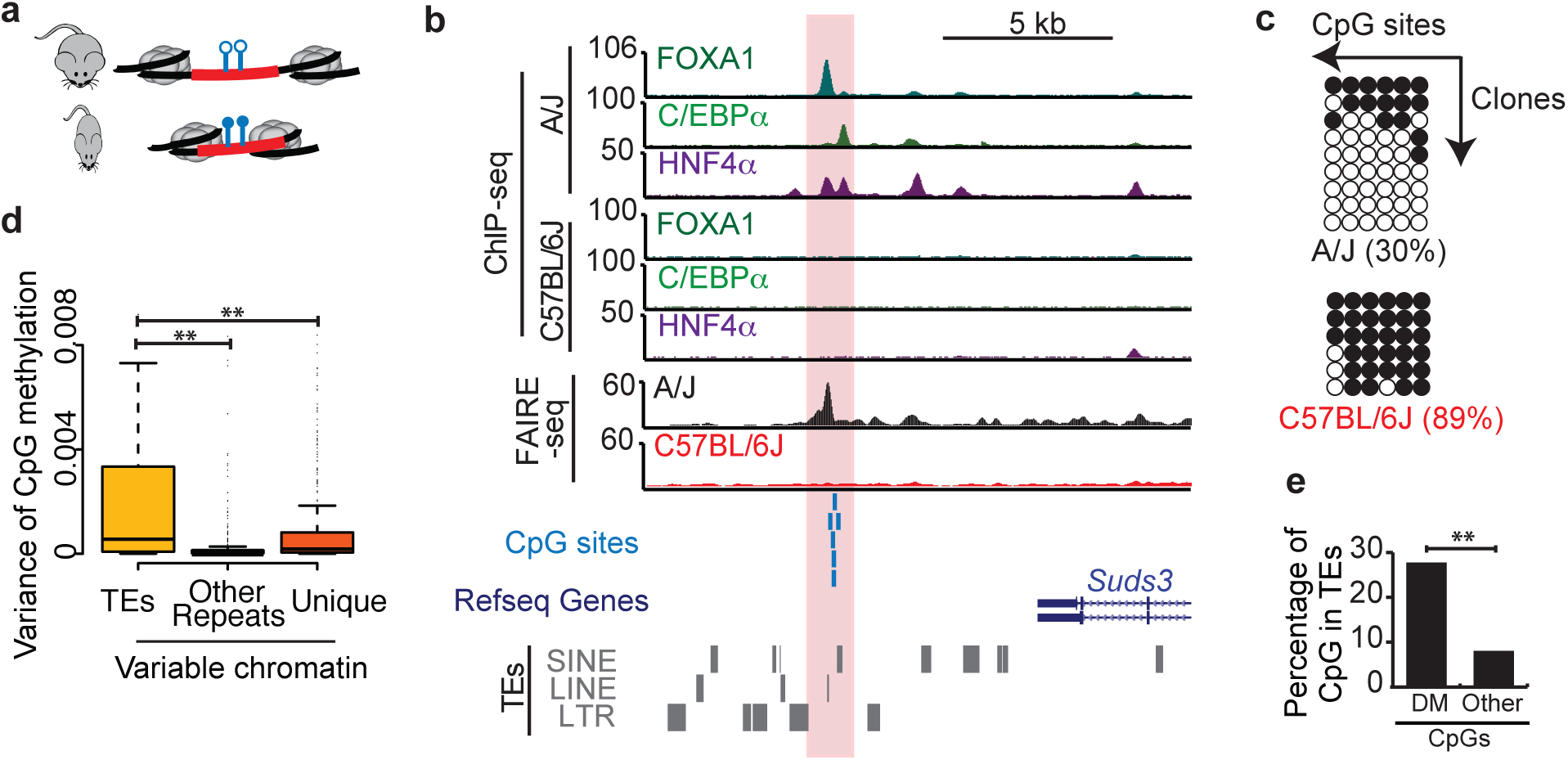
Variable DNA methylation at TEs contributes to chromatin variation. Model of TEs contributing to chromatin variation through differential DNA methylation of TEs. (**b**) Genome browser view of differentially methylated TEs with variable chromatin accessibility. (**c**) Bisulfite sequencing in A/J and C57BL/6J liver of the highlighted region in b. Filled circles represent methylated CpG sites, whereas open circles represent unmethylated sites. (**d**) Boxplot of CpG methylation variance among the seven strains within variable chromatin sites that contain TEs, other repeat elements, or unique sequences. (**e**) Percentage of differential methylated (DM) and other CpGs in 25 inbred strains within TEs (** *p* < 0.001, Fisher’s exact test).

To further validate that the regulatory variation at TEs in liver are not only restricted to the seven inbred strains of mice, we compared the CpG methylation levels from livers of 25 inbred mouse strains [58]. Consistent with our results from the chromatin accessibility variation of seven strains, the differentially methylated (DM) CpG sites among different inbred strains are significantly enriched for TEs compared to other CpG sites (Fig. 7e, *p* < 2.2 × e^−16^, Fisher’s exact test). These results suggest that widespread regulatory variation at TEs is a general feature in mouse liver.

## Discussion

Tissue-specific regulatory regions in the genome are characterized by accessible chromatin [32]. While previous studies have identified a genetic component to chromatin variation [7], the mechanisms underlying the majority of chromatin variation have remained unexplained. We report here that TEs are a major contributor to chromatin variation in liver tissue and furthermore that TE-driven chromatin variation is important for metabolic phenotypes.

We have previously shown that variation in chromatin accessibility across strains of mice depends on genetic factors [30, 39]. We have now extended our study to a total of seven inbred strains that have significant variability in liver phenotypes in response to a HF/HS diet. The variability of the phenotype in these mice resembles the diversity of diet-response in humans [59]. Although in general accessible chromatin sites are less likely to overlap TEs in liver tissue of inbred mice (Additional file 1: Figure S4a, b), we found that chromatin sites with higher variability in different strains are enriched for TEs, specially evolutionarily younger LINEs, as compared to chromatin sites that are common among different strains.

In our study, we used duplicates of each strain of mice for chromatin accessibility profiling and employed a computational pipeline (see Methods) to identify reproducible chromatin variation among different strains of mice. A previous study on C57BL/6J mice showed inter-individual variation of DNA methylation at TEs [60]. We used the 356 regions identified as inter-individual differentially methylated regions [60], and found less than 1% (16/2539) of our variable sites contain inter-individual variability, indicating that the majority of the variable chromatin sites we identified represent sites of variability among different strains of mice.

TEs have been shown to play an important role in expanding the TF binding repertoire during mammalian evolution [24, 27]. Supportive of this, we found that younger and older LINEs have differential chromatin accessibility and are bound by different TFs. Evolutionarily older LINEs are enriched for binding sites of important liver regulatory factors (HNF4α and C/EBPα), indicating their important regulatory roles in liver metabolism. In contrast, younger LINEs are enriched for the binding sites of TFs involved in immune response, such as STATs. The relationship between STAT and TEs is intriguing; a recent study has demonstrated that specific TEs play a functional role in immune pathways in human HeLa cells [21]. It is possible that the variable sites uncovered by our studies also contribute to immune pathways regulated by STAT proteins.

Using genomic enrichment analysis, we demonstrate that TE-associated variable chromatin sites are associated with important liver metabolic genes. One of the classical models of TE contribution to phenotypic diversity is the *agouti viable yellow* (*A*^*vy*^) gene, for which a TE exists upstream of the *agouti* gene [61]. Variation of DNA methylation at this TE regulates the expression of the *agouti* gene and thereby leads to differential coat color and obesity susceptibility [61, 62]. Further experimental validation on the TE-associated variable chromatin sites may lead to the identification of more examples like *A*^*vy*^ gene.

We further investigated possible mechanisms of chromatin variation at TEs. TE polymorphism explains at least 6% of the TE contribution to chromatin variation. Epigenetic mechanisms, such as DNA methylation, have been shown to play an important role in suppression of TE activity in somatic cells [17]. We show here that variation of CpG methylation at TEs contributes to chromatin accessibility variation. DNA hypomethylation at specific TEs has been shown to be associated with enhancer activity [19]. Therefore, these TE-associated chromatin sites may have differential enhancer activity in different strains. Future studies on histone modifications may explain more of the association between TE regulation and chromatin accessibility.

Our finding that TEs contribute to chromatin variation and that they are associated with metabolic loci suggests that the phenotypic diversity observed across the strains is associated with regulatory diversity driven by TEs. Further studies in different tissue types and disease systems may reveal the widespread contribution of TEs to phenotypic diversity.

## Conclusions

In summary, our study has revealed that specific classes of TEs, especially younger LINEs, are a major contributor to chromatin accessibility variation in liver of different inbred strains. We further demonstrate that TEs contribute to tissue-specific regulatory networks and downstream phenotypic diversity.

## Methods

### Animal

Mice were obtained from The Jackson Laboratory and were bred at the University of California, Los Angeles. Male A/J, AKR/J, BALB/cJ, C57BL/6J, C3H/HeJ, CBA/J and DBA/2J mice were maintained on a chow diet (Ralston Purina Company) until 8 weeks of age. Then they were given a high-fat, high-sucrose diet (Research Diets D12266B, 16.8% kcal protein, 51.4% kcal carbohydrate, and 31.8% kcal fat) for 8 weeks. During the feeding period, body fat percentage was tracked as described previously [31]. Mice were then humanely euthanized and livers were harvested. All animal study protocols in this study were approved by the Institutional Care and Use Committee (IACUC) at University of California, Los Angeles and by the Institutional Care and Use Committee (IACUC) at the City of Hope.

### Phenotypic characterization of mice

Animals were measured for body fat mass and lean mass before and after 8 weeks of diet by MRI, as previously described [31]. Liver triglyceride (TG) contents were measured as previously described [34]. Hematoxylin and eosin (H&E) staining and Oil red O staining was performed on liver sections by the Pathology Core at the City of Hope using standard procedures.

### FAIRE-seq and alignment

Formaldehyde-Assisted Isolation of Regulatory Elements (FAIRE) was performed on flash frozen liver tissues from two biological replicates in each strain as previously described [32]. Isolated FAIRE DNA fragment from each sample were barcoded and sequenced on the Illumina HiSeq 2500 to produce 100x100bp paired-end reads.

In order to eliminate the mapping biases caused by inter-strain sequence variation, we first generated pseudo-genome for each non-reference strains by introducing SNPs from each strain into the reference mouse genome (mm9) [36]. We then mapped FAIRE-seq reads from each replicate to the appropriate pseudo-genome using bowtie1 [63] and only reads that could be mapped to single location in the genome were retained. Aligned reads were further filtered to exclude improperly paired reads and PCR duplicates. Overall, we obtained around 17 million uniquely mapped non-duplicate reads in each sample (Additional file 2: Table S1). Wiggle tracks were generated for visualization on the UCSC Genome Browser [64].

For the analysis of FAIRE-seq and RNA-seq coverage at LINEs (Fig. 3), we mapped reads to the reference genome using bowtie2 with the local alignment option [43], as described previously [44]. Unlike bowtie1 with unique mapping mode, the bowtie2 alignment method keeps reads with multiple alignments and reports one of the best alignments [43]. Therefore, the reads from highly similar TE elements can be mapped to a given subfamily of TE.

### Accessible chromatin detection and analysis

To identify accessible chromatin sites from FAIRE-seq reads for each library, F-seq was used with default parameters and a 400bp feature length [65]. To find reproducible peaks across replicates, we utilized the Irreproducible Discovery Rate (IDR) framework [66]. To obtain a union set of accessible chromatin sites from the seven strains, we used the mergeBed function with default parameters [67].

To identify variable chromatin sites among different strains, we first counted the FAIRE-seq reads from each FAIRE-seq library overlapping with the union set of accessible chromatin sites. We normalized the read counts using quantile normalization [68]. We then used DESeq [37] to identify variable chromatin sites among the seven strains, as has been applied previously [7]. We ranked the accessible chromatin sites by adjusted *p*-values from DESeq. The 5% with smallest adjusted *p*-values were considered as variable chromatin sites, whereas the 5% with biggest adjusted *p*-values were considered as least variable (common) chromatin site among the seven strains (Additional file 1: Figure S2). The correlation of FAIRE-seq signal and local sequence variation (Additional file 1: Figure S3) was analyzed as previously described [7].

### Identification of TE-associated chromatin sites

To identify accessible chromatin sites associated with TEs, we used intersectBed [67] to find the accessible chromatin sites that overlap with TEs that annotated by RepeatMasker [38] for the mouse genome (mm9). The age of TEs were calculated as: age = divergence / substitution rate, as previously described [27]. The divergence rates (number of mismatches) for all TEs were obtained from the RepeatMasker annotation file [38]. We used the substitution rates as 4.5 **×**10 ^−9^ per site per year for the mouse genome [11, 27].

### Motif search

To study the potential regulatory roles of LINEs, we used HOMER (version 4.8) motif finding tool (findMotifsGenome.pl) [69] to scan the motifs of known TFs in variable chromatin sites containing younger (< 40 million years (Myrs)) or older LINEs (>= 40 Myrs). Motifs with *p*-value of enrichment less than 0.01 that occurred in more than 10% of the targeted sequences are selected. Highly similar motifs were combined by using joinmotifs tool [70], and only one of the similar motifs is reported. HOMER motif scanning tool (scanMotifGenomeWide.pl) [69] was used to scan occurrence of HNF4α and STAT motif in the whole genome. The putative binding sites were defined by the motif occurrence within accessible chromatin regions identified in C57BL/6J mice liver, similar to what has been reported before [51].

### Bisulfite Sanger sequencing

Genomic DNA from liver tissue was bisulfite-treated according to the manufacturer’s instructions (EpiTect Bisulfite Kit, QIAGEN, USA). Converted genomic DNA was used for PCR reaction (primers: AGGGGTTTTAGGTTTTAGGAAGAG (F), CCAAAACTACTCAAAAAAACCCTATC(R)). Purified PCR products were cloned into pDrive Cloning vector (PCR Cloning Kit, QIAGEN, USA). White colonies were selected through blue/white screening analyzed with Sanger sequencing.

### DNA methylation data

DNA methylation data from reduced representation bisulfite sequencing (RRBS) was obtained from GEO (accession number GSE67507 [58]). Similar to previous analyses [58], only CpG sites were included for analysis. For simplicity, we deleted the small amount of polymorphic CpGs in the seven strains from the methylation data. Differentially methylated (DM) regions were identified as variance bigger than 0.05, and the range of methylation differences bigger than 0.75.

### Gene Ontology (GO) Analysis

In order to investigate the enriched biological function of the genes nearby accessible chromatin sites, we submitted the mouse coordinates (UCSC mm9) of accessible chromatin sites to Genomic Regions Enrichment of Annotations Tool (GREAT) version 3.0.0 [52]. Gene regulatory regions were defined using default parameters (5 kb upstream, 1 kb downstream, and up to 1000 kb distal) and included significant associations for “GO Terms Biological Process”. Only terms that were below a false discovery rate (FDR) of 0.01 were reported.

### RNA-seq and ChIP-seq data

RNA-seq data from livers of C57BL/6J and A/J mice fed with HF/HS diet were obtained from GEO (accession number GSE55581 [30] and GSE75984 [39]). ChIP-seq sites of liver TFs (HNF4~, C/EBPa, and FOXA1) from C57BL/6J and A/J liver tissues are downloaded from ArrayExpress (accession number E-MTAB-1414 [47]). CTCF ChIP-seq sites were obtained from GEO (accession number GSM918715 [4]).

## Declarations

### List of abbreviations

TE: transposable element; FAIRE: Formaldehyde-Assisted Isolation of Regulatory Elements; HF/HS: high-fat, high-sucrose; SNP: single nucleotide polymorphism; LINE: Long interspersed nuclear element; SINE: Short interspersed nuclear element; LTR: Long terminal repeat; TF: transcription factor; TG: triglyceride; H&E: Hematoxylin and eosin; Adi1: acireductone dioxygenase 1; Trappc12: trafficking protein particle complex 12; Myrs: million years; HNF4α: Hepatocyte Nuclear Factor 4, Alpha; CEBPα: CCAAT/Enhancer Binding Protein, Alpha; FOXA1: Forkhead Box A1; STAT: Signal transducers and activators of transcription; GAS: gamma-activated sequence; Enpp1: ectonucleotide pyrophosphatase/phosphodiesterase 1; Suds3: suppressor of defective silencing 3 homolog (S. cerevisiae); DM: differentially methylated; RRBS: reduced representation bisulfite sequencing.

### Ethics approval and consent to participate

All animal study protocols in this study were approved by the Institutional Care and Use Committee (IACUC) at University of California, Los Angeles and by the Institutional Care and Use Committee (IACUC) at the City of Hope.

### Availability of data and material

The sequence data have been deposited in the NCBI GEO repository (GSE75770).

### Authors’ contributions

JD, AL and DES designed the study. JD, CT and BWP performed the experiments. JD, AL and DES conducted the analysis. JD, AJL, RN and DES prepared the manuscript. All authors read and approved the final manuscript.

## Acknowledgements

We would like to acknowledge other members of the Schones and Lusis laboratories for helpful discussions and comments. We thank Sadhan Das, Xiaoxiao Ma, and Kaniel Cassady for suggestions on the manuscript.

